# Convergent Amino-Acid Substitutions Link Primate Longevity to Male-Fertility Genes

**DOI:** 10.64898/2026.09.04.749475

**Authors:** Fabio Barteri, Noelia Rodriguez-Perez, Miguel Ramon, Eva Brigos, David Juan, Gerard Muntané, Arcadi Navarro

**Affiliations:** Institute of Evolutionary Biology (IBE, UPF–CSIC), Department of Medicine and Life Sciences, Universitat Pompeu Fabra, Parc de Recerca Biomèdica de Barcelona (PRBB), Barcelona, Spain; Systems Biology Department, Spanish National Center for Biotechnology (CNB-CSIC), Madrid, Spain; Hospital Universitari Institut Pere Mata, Institut de Recerca Biomèdica Catalunya Sud, Universitat Rovira i Virgili, Reus, Spain; Centro de Investigación Biomédica en Red en Salud Mental (CIBERSAM), Spain; BarcelonaBeta Brain Research Center, Pasqual Maragall Foundation, Barcelona, Spain; Institució Catalana de Recerca i Estudis Avançats (ICREA) and Universitat Pompeu Fabra, Barcelona, Spain; Center for Genomic Regulation (CRG), The Barcelona Institute of Science and Technology, Barcelona, Spain

## Abstract

Maximum lifespan varies widely among mammals, but the genomic basis of this variation remains incompletely understood. We used body-mass-corrected longevity and a phylogenetically informed phenotype-shift framework to compare 126 primate species, selecting seven long-lived and six short-lived species for convergent amino-acid substitution analysis. Stringent pooling across species identified 1,068 recurrent phenotype-associated positions in 934 genes. These genes showed significant agreement with mammal-wide longevity candidates, with 226 genes shared compared with 120 expected by chance (1.88-fold enrichment; P = 1.16 × 10^−22^). Functional analysis detected 17 significant terms, with the strongest coherent signal involving male infertility and decreased male fertility. Our results show that focused primate sampling recovers a non-random component of the broader mammalian longevity signal and point to male reproductive biology as a promising direction for investigating the evolution of lifespan.

## 1 Introduction

The diverse and often unpredictable forces governing lifespan have been debated since antiquity. Nevertheless, maximum lifespan (MLS) is a species-specific trait with a strong phylogenetic signal and an approximately 60-fold range among mammals, indicating substantial genetic determination alongside ecology and life history (de Magalhães and Toussaint, 2002; Ma and Gladyshev, 2017; Farré et al., 2021).

Because lifespan covaries with body size, comparative studies use the longevity quotient (LQ), an allometric residual expressed as observed MLS divided by its model-based expectation. Alternative corrections use mammal-wide or taxon-specific body-mass relationships and, in some formulations, metabolic rate (Austad and Fischer, 1991; Speakman, 2005; de Magalhães et al., 2007; Farré et al., 2021). Cross-species studies connect longevity with genome maintenance, including enhanced DNA repair and reduced accumulation of somatic mutations (Cagan et al., 2022; Firsanov et al., 2025). Longevity also intersects reproduction: human genetic data support antagonistic pleiotropy, with fertility-enhancing alleles associated with reduced longevity (Brigos-Barril et al., 2026).

Using LQ and phylogenetically informed comparative genomics, Farré et al. (2021) identified 2,737 convergent amino-acid substitutions (CAAS) in 2,004 genes distinguishing long- and short-lived mammals, including 1,157 substitutions significantly associated with MLS. Here, we revisit this strategy within primates, leveraging the genomic and phylogenetic catalog of 233 species generated by Kuderna et al. (2023). Our primate-focused CAAS set significantly overlaps the mammalian discoveries of Farré et al. (2021) and shows multiple functional enrichments, with the strongest coherent signal involving male-fertility annotations (Supplementary Methods S1–S4).

## 2 Results and Discussion

### 2.1 Allometrically corrected longevity is phylogenetically structured across primates

The fixed mammal-wide relationship between adult body mass and maximum lifespan was applied to 126 primate species matched to the Kuderna S4 phylogeny, producing a longevity quotient (LQ) that quantified species-level deviations from the allometric expectation (Fig. 1A). LQ retained significant phylogenetic structure across this dataset: Blomberg’s K was 0.146 (P = 3.00 × 10^−4^) and Pagel’s λ was 0.786 (P = 3.94 × 10^−9^; Fig. 1D). Consistent with this structure, an Ornstein–Uhlenbeck model was favoured over Brownian motion for phenotype-shift score (PSS) construction (AIC = −28.83 versus 11.88). The combination of substantial variation around the allometric expectation and detectable phylogenetic dependence provided the basis for identifying repeated longevity contrasts rather than treating species as independent observations. Thus, correcting for body mass did not remove the contribution of shared ancestry to primate longevity, supporting an explicitly phylogenetic comparison (Supplementary Methods S1–S2).

**Figure 1.**
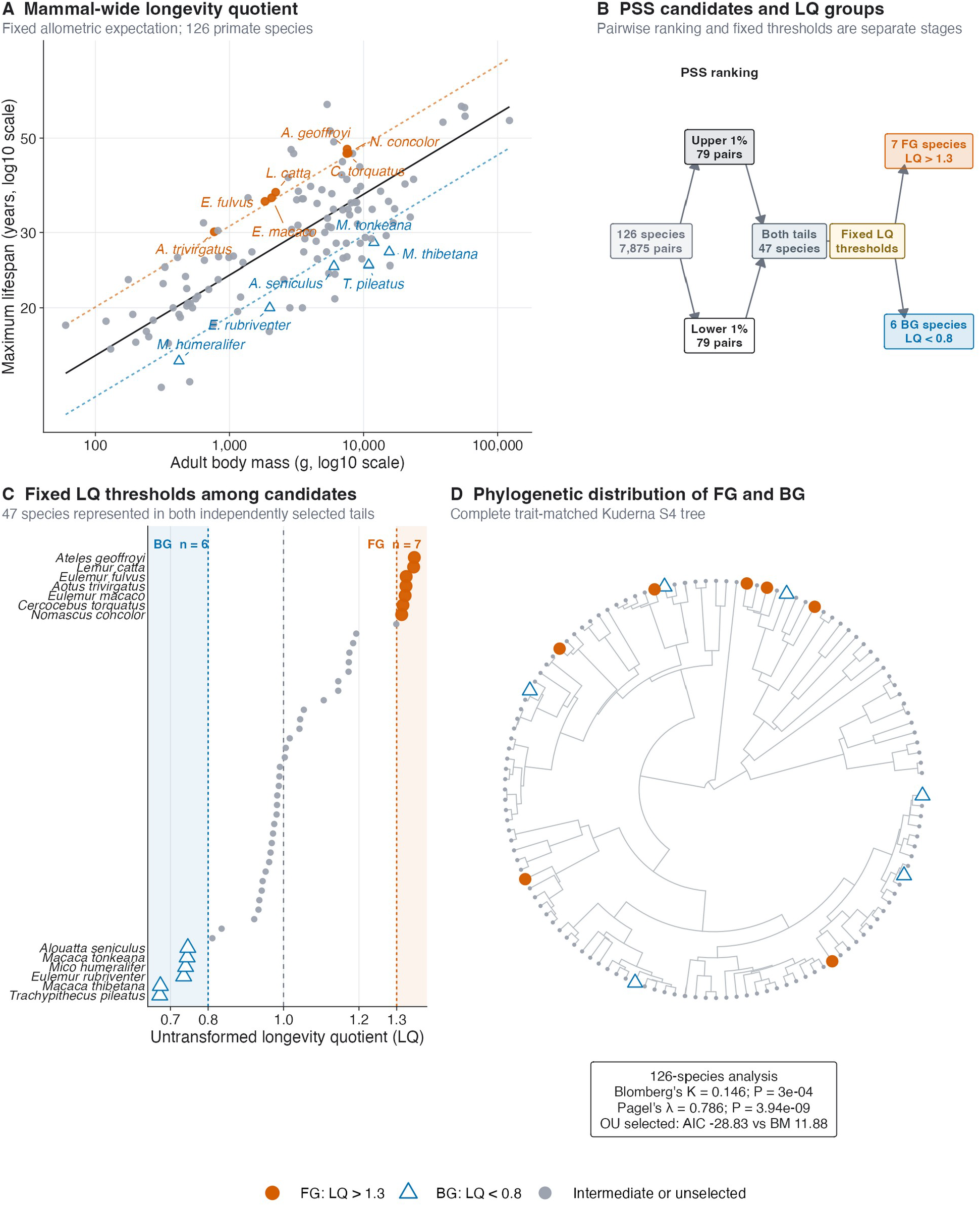
Allometrically corrected longevity identifies repeated phenotypic extremes across the primate phylogeny. Maximum lifespan, body mass, and phylogenetic information were integrated to define the foreground (FG) and background (BG) species used in molecular analyses. **A**, Observed maximum lifespan plotted against adult body mass for the 126 species retained after exact matching to the Kuderna S4 phylogeny. Both axes are logarithmic. The solid black line is the fixed mammal-wide expectation, expected maximum lifespan = 6.47 × body mass (g)^0·189^; dashed lines mark longevity quotient (LQ) values of 0.8 and 1.3. FG species are vermillion circles, BG species are blue triangles, and other species are grey points. **B**, Schematic of candidate construction. Phenotype shift scores (PSSs) ranked all 7,875 unordered species pairs. The independently selected Upper 1% and Lower 1% contained 79 pairs each, and 47 species occurred in both pair-derived species sets. Fixed phenotype thresholds then identified seven FG species with LQ > 1.3 and six BG species with LQ < 0.8. PSS selection and LQ thresholding are separate stages. **C**, Absolute, untransformed LQ for all 47 shared-tail species. Every point represents one species; only the 13 selected species are named. The dashed grey line marks the mammal-wide expectation of LQ = 1. The two coloured dashed lines mark the strict operational thresholds; species with 0.8 ≤ LQ ≤ 1.3 remain unselected. **D**, Placement of the selected species on the complete 126-tip trait-matched phylogeny. The same colour and shape encodings are used throughout. Untransformed LQ retained significant phylogenetic signal (Blomberg’s K = 0.146, 9,999 tip-label permutations, P = 3.00 × 10−4; Pagel’s λ = 0.786, likelihood-ratio test against λ = 0, P = 3.94 × 10−9). The PSS analysis selected an Ornstein–Uhlenbeck model (AIC = −28.83) over Brownian motion (AIC = 11.88). The tree describes the distribution of observed species-level phenotypes and does not infer discrete ancestral longevity regimes.

### 2.2 Phenotype-shift ranking and fixed LQ thresholds identify recurrent longevity extremes

PSS ranked all 7,875 unordered species pairs according to their phylogenetically normalized phenotype differences (Fig. 1B). Independent selection of the upper and lower 1% tails retained 79 pairs in each tail, and 47 species occurred in both pair-derived species sets. Applying fixed absolute thresholds only after this PSS selection identified seven foreground (FG) species with LQ > 1.3 and six background (BG) species with LQ < 0.8 (Fig. 1B,C). These 13 species occupied several separated regions of the complete 126-tip phylogeny rather than a single terminal clade (Fig. 1D). Thus, PSS ranking and absolute LQ thresholding contributed distinct selection stages and yielded phylogenetically distributed phenotype groups for the molecular comparisons. Their distribution limits domination by a single terminal radiation, although the operational PSS tails and LQ cut-offs identify comparative candidates rather than independently inferred longevity regimes (Supplementary Method S2).

### 2.3 Pooled CAAS analysis identifies recurrent foreground-concordant amino-acid changes

Phenotype-consistent FG–BG comparisons were pooled by gene and aligned amino-acid position, and repeated appearances of a species across hypotheses were collapsed before its support was counted (Fig. 2A). The nominal positional P ≤ 0.05 universe comprised 129,622 pooled events in 10,795 genes (Fig. 2C). Requiring support from at least five unique FG and five unique BG species reduced this universe to 2,984 events in 2,104 genes. Complete FG amino-acid concordance and retention of Patterns 1 and 2 then produced the strict primary set of 1,068 unique positions in 934 genes. Pattern 1, in which FG and BG species each carried a different shared state, accounted for 600 positions (56.2%); Pattern 2, in which the shared FG state was opposed by heterogeneous BG states without a BG dominance filter, accounted for 468 positions (43.8%; Fig. 2B,D). Most strict events were supported by five species in each phenotype group, whereas allowing FG dominant frequency ≥0.80 yielded a broader sensitivity set of 1,713 positions in 1,397 genes rather than an intermediate stage of the strict analysis (Fig. 2C,D). The sharp reduction across filters shows that recurrent species support and foreground concordance, rather than nominal positional significance alone, drove the primary set. These candidates therefore represent repeated phenotype-associated substitutions, not experimentally validated determinants of lifespan (Supplementary Method S3).

**Figure 2.**
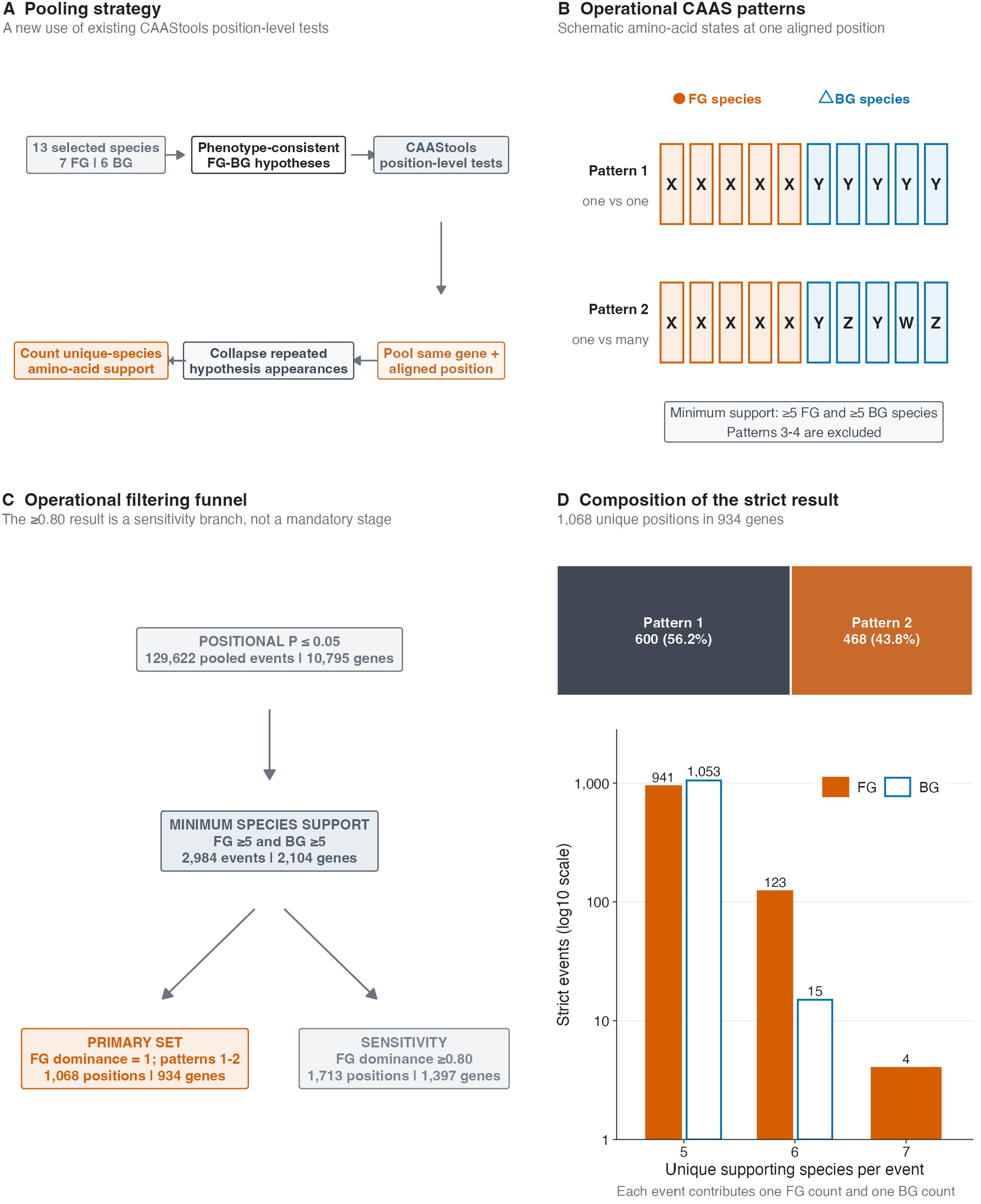
Pooling phenotype-consistent CAAS comparisons identifies recurrent foreground-concordant amino-acid changes. A pooled application of existing CAAStools tests was used to prioritize convergent amino-acid substitutions (CAAS) supported across multiple phenotype-consistent primate comparisons. **A**, Analysis workflow. The 13 species defined in Figure 1 (seven foreground [FG] and six background [BG] species) generated multiple phenotype-consistent FG-BG hypotheses. CAAStools position-level tests were pooled by gene and aligned amino-acid position. Repeated appearances of a species across hypotheses were collapsed before support was counted, so hypotheses were not treated as independent biological replicates. **B**, Operational definitions of the two patterns retained in the strict analysis. Pattern 1 contains one FG amino acid and one non-overlapping BG amino acid. Pattern 2 contains one FG amino acid and multiple non-overlapping BG amino acids. Letters denote schematic amino-acid states rather than selected genes. Both groups required support from at least five unique species. Patterns 3 and 4, which contain multiple FG amino acids, were excluded from the strict complete-FG-concordance set. **C**, Filtering funnel. The positional nominal *P* ≤ 0.05 universe contained 129,622 pooled events in 10,795 genes. Requiring support from at least five FG and five BG species retained 2,984 events in 2,104 genes. Complete FG amino-acid concordance (dominant frequency = 1) produced the primary set of 1,068 unique positions in 934 genes. The 1,713-position, 1,397-gene result obtained at FG dominant frequency ≥ 0.80 is shown as a sensitivity branch rather than a mandatory stage of the primary analysis. No BG amino-acid dominance filter was applied. **D**, Composition of the strict primary result. Pattern 1 accounted for 600 positions (56.2%) and Pattern 2 for 468 positions (43.8%). The lower plot shows the number of strict events supported by five, six, or seven FG species and by five or six BG species; the vertical axis is logarithmic. Every event contributes one FG support count and one BG support count. All thresholds are operational analysis filters rather than universal biological cutoffs.

### 2.4 Strict CAAS genes show significant agreement with the Farré et al. discovery sets

The 934 strict convergent amino-acid substitution (CAAS) genes were compared with the discovery sets of Farré et al. (2021) using the 16,133 genes represented in the alignment as the statistical universe (Fig. 3A). Across all discovery scenarios, 226 genes overlapped compared with 120.0 expected by chance, corresponding to a 1.88-fold excess (one-sided hypergeometric P = 1.16 × 10^−22^). Excess overlap was independently observed for discovery scenarios 1–2 (202 versus 104.2 expected; 1.94-fold; P = 1.44 × 10^−21^) and scenario 3 (54 versus 25.4 expected; 2.13-fold; P = 1.16 × 10^−7^). The agreement across scenario classes indicates that the primate-focused analysis recovered a non-random component of the mammal-wide longevity signal, despite its denser sampling of a single order. However, this agreement was evaluated at gene level and does not imply recurrence of the same amino-acid position or substitution (Supplementary Method S4).

**Figure 3.**
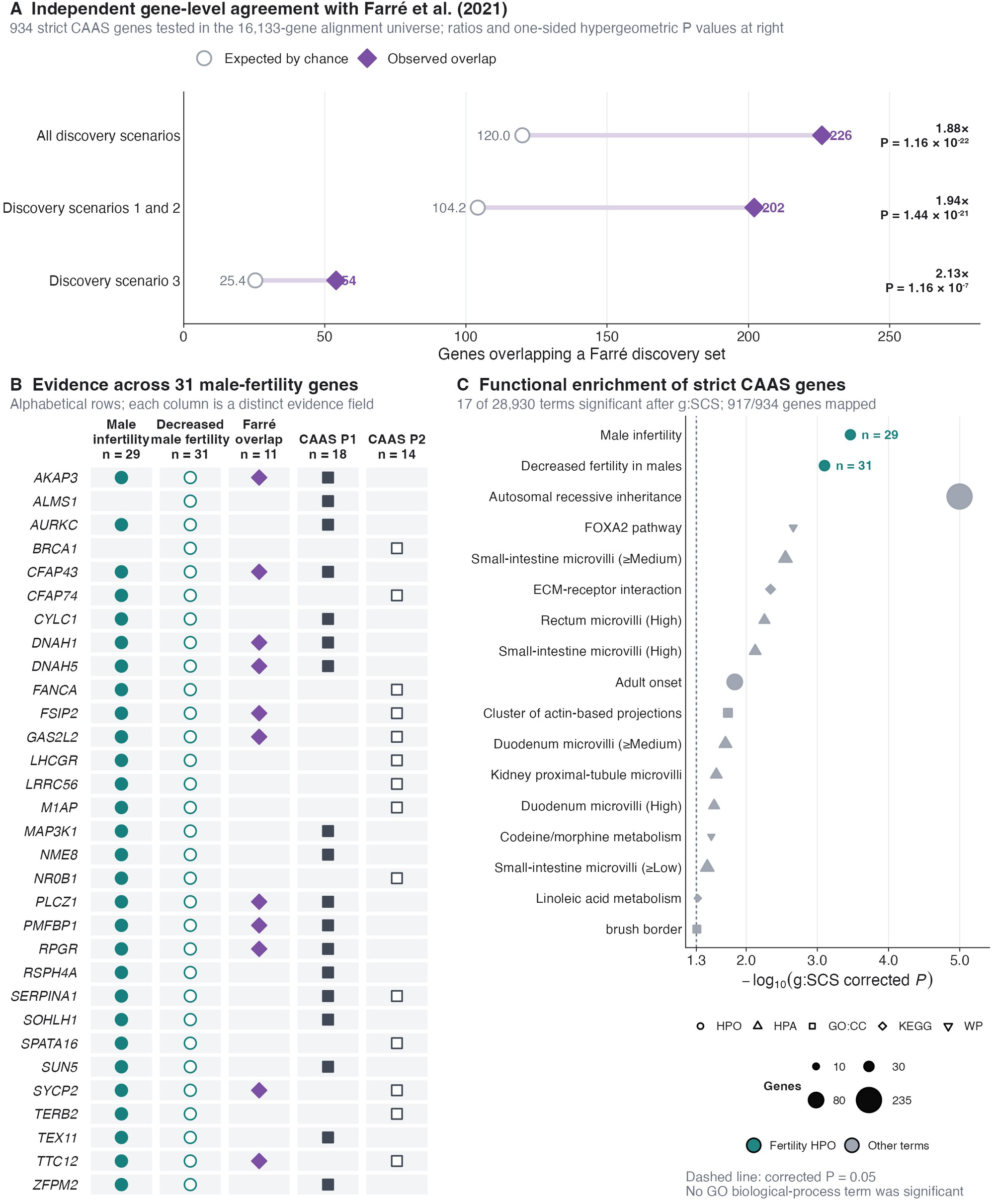
Independent gene-level evidence links longevity-associated molecular evolution to male fertility. The strict convergent amino-acid substitution (CAAS) gene set was evaluated against an independent primate-aging study and functionally characterized. **A**, Gene-level overlap between the 934 strict CAAS genes and the discovery sets reported by Farré *et al*. (2021). Across all discovery scenarios, 226 genes overlapped, compared with 120.0 expected by chance (observed/expected = 1.88; one-sided hypergeometric *P* = 1.16 × 10^−22^). Discovery scenarios 1 and 2 contained 202 overlapping genes versus 104.2 expected (1.94-fold; *P* = 1.44 × 10^−21^), whereas discovery scenario 3 contained 54 versus 25.4 expected (2.13-fold; *P* = 1.16 × 10^−7^). Tests used the 16,133 genes represented in the alignment as the universe. Agreement is evaluated at gene level and does not imply recurrence of the same amino-acid position or substitution. **B**, Evidence matrix for the union of the two significant male-fertility Human Phenotype Ontology (HPO) terms. Of 31 genes annotated to decreased fertility in males, 29 were also annotated to male infertility; 11 overlapped a Farré discovery set, 18 contained a strict Pattern-1 CAAS, and 14 contained a strict Pattern-2 CAAS. One gene contained both strict CAAS patterns. Rows are alphabetical. Filled and outlined symbols distinguish related evidence fields; absence of a symbol means that the corresponding evidence was not present in the frozen source tables. HPO annotations are functional annotations and are not direct observations of fertility in the primate species analysed. **C**, All 17 terms significant among 28,930 tested terms after g:SCS multiple-testing correction. Enrichment used 917 mapped genes from the 934-gene strict set and an effective domain of 15,965 genes. The horizontal coordinate is −log_10_ of the corrected *P* value, point area represents the number of intersecting genes, and point shape denotes the source database. Male infertility (29 genes; corrected *P* = 3.43 × 10^−4^) and decreased fertility in males (31 genes; corrected *P* = 7.94 × 10^−4^) are highlighted. Other significant terms include microvillar and actin-based cell-projection annotations and extracellular-matrix receptor interaction. The dashed line marks corrected *P* = 0.05. No Gene Ontology biological-process term was significant. HPA, Human Protein Atlas; HPO, Human Phenotype Ontology; WP, WikiPathways.

### 2.5 Functional enrichment links longevity-associated CAAS genes to male fertility

Of the 934 strict CAAS genes, 917 mapped in the functional analysis, which identified 17 significant terms among 28,930 tested terms after g:SCS correction (Fig. 3C). The leading phenotype annotations included male infertility (29 genes; corrected P = 3.43 × 10^−4^) and decreased fertility in males (31 genes; corrected P = 7.94 × 10^−4^). Within the 31-gene fertility union, 11 genes also overlapped a Farré discovery set, 18 contained a strict Pattern-1 CAAS, and 14 contained a strict Pattern-2 CAAS; one gene contained both patterns (Fig. 3B). Other significant terms involved microvilli, actin-based cell projections, and extracellular-matrix receptor interaction, whereas no Gene Ontology Biological Process term was significant (Fig. 3C). The male-fertility signal is notable in light of human genetic evidence for antagonistic pleiotropy between reproduction and longevity, in which alleles associated with greater reproductive success tend to predict shorter lifespan (Brigos-Barril et al., 2026). This correspondence makes reproductive and male-germline biology plausible components of the longevity-associated genomic landscape.

Nevertheless, the Human Phenotype Ontology memberships are annotation-based and neither demonstrate altered fertility in the primates analysed nor establish that a reproductive trade-off causally mediates their longevity differences (Supplementary Method S4).

## 3 Conclusions

The comparative analysis of lifespan variation across primates identified recurrent phenotype-associated amino-acid substitutions in genes that significantly overlap with mammal-wide longevity candidates. This agreement indicates that the primate-focused analysis recovers part of a broader genomic signal associated with lifespan evolution. Functional enrichment further pointed towards male-fertility phenotypes, placing reproductive biology among the most prominent themes emerging from the data. Although this signal does not establish a causal relationship between fertility and longevity, it identifies a specific and biologically plausible direction for future research. Broader comparative analyses and functional investigation of the implicated genes will be required to determine whether, and through which mechanisms, male reproductive biology contributes to the evolution of longevity.

## 4 Methods

### 4.1 Phenotype data and phylogeny

Maximum lifespan (trait PTD00065, https://pgarchive.github.io) was obtained from the non-human-primate phenomic dataset, which incorporates AnAge records (de Magalhães and Costa 2009), and adult body mass (PTD00014) from Galán-Acedo et al. (2019). Analyses retained species with finite, positive values for both traits. Body mass was converted from kilograms to grams and expected maximum lifespan was calculated with the fixed mammalian allometry of de Magalhães et al. (2007): expected lifespan = 6.47 × mass^0·189^. The longevity quotient (LQ) was observed divided by expected lifespan; untransformed LQ was used. Species names were matched to the supplementary phylogeny of Kuderna et al. (2023), which was pruned to the 126 matched species (Supplementary Method S1).

### 4.2 Phenotypic contrasts and phylogenetic signal

Brownian-motion and Ornstein–Uhlenbeck models (Hansen 1997) were fitted to LQ with geiger::fitContinuous (Pennell et al. 2014); the OU model was selected only when its AIC was at least two units lower. Pairwise phenotype-shift scores (PSS; Barteri et al. 2026) were calculated for all 7,875 unordered species pairs under the selected model. Pairs were ranked by PSS, and the upper and lower 1% (79 pairs per tail) were retained. Among species represented in both tails, foreground and background candidates were defined a priori as LQ > 1.3 and LQ < 0.8, respectively; the tails were used descriptively rather than as significance tests. Phylogenetic signal in LQ was quantified using Blomberg’s K with 9,999 tip-label permutations (Blomberg et al. 2003) and Pagel’s maximum-likelihood λ, tested against λ = 0 by a likelihood-ratio test (Pagel 1999), using phytools::phylosig (Revell 2012) (Supplementary Method S2).

### 4.3 Pooled discovery of convergent amino-acid substitutions

Convergent amino-acid substitutions were screened across 16,133 protein alignments using a locally extended pooled-discovery implementation of CAAStools (Barteri et al. 2023). The seven foreground and six background species yielded 525 possible 4-versus-4 hypotheses; 100 were selected deterministically using a seeded SHA-256 procedure (seed 260811). For each alignment and hypothesis, patterns 1–3 were evaluated at positions represented by at least three non-gap sequences per group and with a maximum gap fraction of 0.5. A positional hypergeometric test provided nominal P values. Compatible calls were pooled by gene, alignment position, and amino-acid signature; repeated appearances of a species across hypotheses were collapsed so that each species contributed once per event. Primary candidates required positional P ≤ 0.05, at least five unique species in each group, fixation of the dominant foreground residue, and CAAStools pattern 1 or 2. The workflow was orchestrated with Nextflow (Di Tommaso et al. 2017) (Supplementary Method S3).

### 4.4 Functional enrichment and external validation

Functional enrichment was performed with the g:Profiler g:GOSt API (Reimand et al. 2007; Kolberg et al. 2023), querying human orthologues against a custom background comprising all 16,133 screened genes. All annotation sources and electronically inferred Gene Ontology annotations were included; significance used the g:SCS correction at 0.05. The API was queried on 26 August 2026. Gene-level overlap with the complete convergent-substitution set of Farré et al. (2021) was evaluated by one-sided hypergeometric tests using the same 16,133-gene universe, both across all reported scenarios and separately for scenarios 1–2 and 3. A sensitivity analysis repeated downstream summaries after excluding cilia- and flagella-associated candidates. Analyses used R 4.3.1; seeds, configurations, intermediate tables, and software versions were retained for reproducibility (Supplementary Method S4).

## CRediT Authorship Contribution Statement

Fabio Barteri: Project administration; Supervision.

Noelia Rodriguez-Perez: Formal analysis; Validation.

Miguel Ramon: Formal analysis; Validation.

Eva Brigos: Formal analysis; Validation.

David Juan: Supervision; Writing – review & editing.

Gerard Muntané: Supervision; Writing – review & editing.

Arcadi Navarro: Conceptualization; Supervision.

