## Supplementary Notes for "Convergent Amino-Acid Substitutions Link Primate Longevity to Male-Fertility Genes"

### Supplementary Methods

#### Supplementary Method S1. Phenotype data, taxonomic reconciliation, and longevity quotient

Phenotypic records were obtained from the frozen non-human-primate phenomic dataset `nhp.phenomic.dataset.csv`, which contained 526 unique `GroupName` records. Each identifier encoded superfamily, family, genus, and species as four underscore-delimited fields. Maximum lifespan (MLS) in years was taken from trait `PTD00065` (`Maximum.lifespan..y.`); its local metadata attribute the observations to the Van Schaik/Isler compilation and AnAge. Adult body mass in kilograms was taken from trait `PTD00014` (`BodyMass_kg`) and was attributed to Galán-Acedo et al. (2019). The available source metadata did not contain record-level sex, population, captive/wild status, sample size, uncertainty, or access date. Each row already represented one unique species record; no sex- or population-specific observations were therefore collapsed or averaged.

Trait values were converted to numeric form, and body mass was multiplied by 1,000 to obtain grams. A record was eligible only when both MLS and body mass were finite and strictly positive. Of the 526 records, 162 met these criteria and 364 were excluded: 303 lacked valid MLS only, one lacked valid body mass only, and 60 lacked both. All records, including exclusions, were retained in an audit table with their original and standardized values, trait identifiers, source fields, inclusion status, and explicit exclusion reason. Genomic sample counts were added as annotations but were not used to determine phenotype eligibility.

Species names were constructed directly as `Genus_species`. Downstream analyses removed apostrophes and replaced spaces with underscores, rejected duplicate species identifiers, and matched names by exact string identity to the terminal labels of the Kuderna et al. (2023) Data S4 phylogeny. No approximate matching, synonym substitution, or undocumented manual correction was applied. Thirty-six of the 162 phenotype-eligible species lacked an exact tree match, yielding the same 126-species set for phylogenetic-signal estimation, evolutionary-model fitting, and pairwise phenotype-shift analysis. The complete tree contained 236 tips; 110 tips lacked an eligible LQ observation and were removed when the tree was pruned.

Expected MLS was calculated from the fixed mammal-wide allometry of de Magalhães et al. (2007), without re-estimating either coefficient in primates:

$$\textit{expected MLS (years)} = 6.47 \times \textit{body mass (g)}^{0.189}; \quad \textit{LQ} = \textit{observed MLS} / \textit{expected MLS}$$

The computation was performed on the natural-log scale and exponentiated. An LQ of 1 denotes the lifespan expected under this mammal-wide relationship; it is not a significance boundary or a direct measure of ageing rate. Both LQ and log(LQ) were archived, but untransformed LQ was used for all primary thresholding, phylogenetic-signal, and phenotype-shift analyses. Phenotype attrition, observed-versus-expected MLS, the complete LQ distribution, and family-level coverage are shown in supplementary figure S1; the full record audit and 126-species analysis table are provided in supplementary table S1 and supplementary data S1.

#### Supplementary Method S2. Phylogenetic modelling and phenotype-shift selection

The Kuderna et al. (2023) Data S4 tree was rooted, fully bifurcating, ultrametric, and supplied with finite positive branch lengths. It was pruned to the 126 exactly matched species without modifying the retained topology or branch lengths. Brownian-motion (BM) and single-optimum Ornstein–Uhlenbeck (OU) models were fitted by maximum likelihood to untransformed LQ with `geiger::fitContinuous`, one computational core, and the package's default optimization settings and parameter bounds (Pennell et al. 2014). Model selection preceded score construction. The prespecified rule selected OU only when `AIC_OU + 2 < AIC_BM`; otherwise BM was retained. The fitted BM model had $\sigma^2 = 0.00752788$ and AIC 11.88333, whereas the OU model had $\sigma^2 = 0.01384447$, $\alpha = 0.11520237$, and AIC −28.83019. OU was therefore used to construct the covariance matrix underlying the phenotype-shift score (PSS). This model choice describes relative statistical support and was not treated as evidence for discrete adaptive regimes or stabilizing selection on longevity.

Phylogenetic signal was estimated independently with `phytools::phylosig` (Revell 2012). Blomberg's K was evaluated against a null distribution generated by 9,999 random permutations of the observed LQ values among the 126 fixed tree tips, using seed `20260817` (Blomberg et al. 2003). The implementation reports the upper-tail randomization probability that permuted tip assignments produce a signal statistic at least as extreme in the tested direction. Pagel's $\lambda$ was estimated by maximum likelihood within the bounds implemented by `phylosig`, and evidence against phylogenetic independence was assessed by the likelihood-ratio comparison of the fitted model with the constrained null $\lambda = 0$ (Pagel 1999). Both tests used the same tip order and untransformed trait vector.

PSS was calculated for all C(126,2) = 7,875 unique unordered species pairs following Barteri et al. (2026). For species i and j, the absolute difference was $\Delta_{ij} = |x_i - x_j|$, and its OU-based variance was $V_{ij} = C_{ii} + C_{jj} - 2C_{ij}$. The standardized contrast $z_{ij} = \Delta_{ij} / \sqrt{V_{ij}}$ was converted to the central-normal component $S_{ij} = 2\Phi(z_{ij}) - 1$. The ranked score was

$$PSS_{i,j} = S_{i,j} \times [(\Delta_{i,j} / max(\Delta)) / (d_{i,j} / max(d))]$$

where $d_{ij}$ is patristic distance and both maxima were taken over the complete pair universe. Pair generation followed tree-tip order; missing and non-finite values and duplicate species labels were rejected before calculation. Rows were sorted in descending order of the `FinalScore` column. No exact score ties occurred, so tail membership did not require an additional tie-breaking rule.

The upper and lower 1% tails were selected independently as rank-based descriptive subsets. Their common size was ceiling(7,875 × 0.01) = 79 pairs. Species represented in either tail were extracted, and the intersection of the two pair-derived species sets contained 47 taxa. Fixed LQ thresholds were applied only within this shared-tail set: foreground (FG) required LQ > 1.3, background (BG) required LQ < 0.8, and values at either boundary or between them were unselected. This yielded seven FG species, six BG species, and 34 intermediate shared-tail candidates. Tail participation was a screening condition rather than a significance test, repeated pairs sharing a species were not biological replicates, and the FG/BG labels were not inferred evolutionary regimes. No alternative PSS-tail or LQ-threshold grid was frozen; consequently, no claim of threshold stability is made. The complete ranked distribution and the two selection stages are documented in supplementary figure S2, with model and signal summaries in supplementary table S2, species membership in supplementary table S3, and all pairwise scores in supplementary data S2.

#### Supplementary Method S3. Pooled discovery and quality control of convergent amino-acid substitutions

The molecular screen used 16,133 gene-level protein alignments in relaxed PHYLIP format (`*.filter2.phy`) and the seven FG and six BG species defined above. Each alignment was treated as one screened gene unit. Where an event table was non-empty, its CAAStools `gene` field was authoritative; for valid header-only results the same basename-parsing rule used by CAAStools recovered the identifier. This procedure retained alignments whose reference name was non-human and filenames containing internal dots. No additional paralogy or transcript reassignment was inferred downstream of the frozen alignment collection. Gaps were represented by `-`; absent species were recorded as missing, and only present non-gap amino acids contributed to group support. Ambiguous amino-acid characters were retained as observed states rather than imputed. CAAStools alignment coordinates are zero-based column indices; they are not automatically equivalent to one-based ungapped human-protein residue numbers.

Every hypothesis sampled four distinct species without replacement from each phenotype group. The full design comprised C(7,4)C(6,4) = 525 unique 4-FG-versus-4-BG configurations. A prespecified subset of 100 limited the number of evaluations per alignment while repeatedly representing every selected species; it was not treated as 100 independent evolutionary replicates. Candidate configurations were constructed in the fixed order of the complete FG and BG pool files, encoded as tab-separated seed, FG subset, and BG subset strings, ranked by their SHA-256 digest, and the first 100 were retained using seed `260811`. The realized manifest had SHA-256 `46c47c2566a8ccb7917102e830e6f9067d3e76bec72110fb1b858d20a1fcadde` and was generated once at run level, then reused for every alignment. FG species occurred in 50–67 configurations and BG species in 64–70 configurations. Validation required four unique eligible FG and four unique eligible BG species in every row. The exact configurations and participation counts are reported in supplementary table S4 and its machine-readable manifest.

A locally extended version of CAAStools 0.9 beta (Barteri et al. 2023) was run in `pooled-discovery` mode. Within each 4-versus-4 hypothesis, a position was evaluable when at least three present non-gap sequences were available in each group. General per-group missing-species and gap-count ceilings were disabled, while positions with a gap fraction greater than 0.5 across the alignment were excluded. CAAStools patterns 1–3 were evaluated at this stage. The local extension attached an upper-tail hypergeometric probability to the complete discovery result at each alignment position, using group sizes implied by the hypothesis design, and retained positions with nominal `positional_pvalue <= 0.05`. This value differs from `event_nominal_pvalue`, which is calculated conditionally after an amino-acid event has been inferred. Only `positional_pvalue` was used in the primary discovery filter; neither probability was corrected experiment-wide.

The production workflow was orchestrated with Nextflow DSL2 (Di Tommaso et al. 2017) and executed through SLURM as run `run-260819161121`. One shared process generated the hypothesis manifest, after which one job per alignment used one CPU, 2 GB RAM, and a 30-min limit on the `std-cpu` partition, with at most 100 queued submissions and a submission rate of ten jobs per second. Failed or timed-out gene tasks were configured not to stop the remaining run, whereas failure of hypothesis preparation was fatal. The archived analytical input contained all 16,133 per-gene event tables: 10,795 contained at least one retained event and 5,338 were valid header-only files. Thus, no result table was missing from the downstream universe. The environment used OpenJDK 17, Python 3.11, Biopython, DendroPy, NumPy, and SciPy in the `phyloq` Conda environment.

Positive hypotheses were not counted directly as evidence. Calls at the same gene and zero-based alignment position were merged when their FG and BG amino-acid signatures were compatible; incompatible signatures remained separate event rows. The pooled table retained every contributing hypothesis identifier and designated one primary event per position. Within an event, each species was counted at most once regardless of how many positive hypotheses contained it. `fg_support_count` and `bg_support_count` therefore refer to unique species, and dominant-amino-acid frequencies use the corresponding unique supporting species as denominators. Pattern 1 denotes one shared FG state opposed by one different shared BG state; Pattern 2 denotes one shared FG state opposed by heterogeneous BG states; Pattern 3 reverses that asymmetry; and Pattern 4 contains multiple states on both sides. Patterns 3 and 4 do not satisfy complete FG concordance and were excluded from the strict primary endpoint.

Per-gene event tables were processed in sorted filename order, required to share a schema, and merged without modifying the raw outputs. The broad nominal universe comprised 129,622 event rows, 126,598 unique gene-position combinations, and 10,795 genes. For descriptive support, balanced support was defined as

$$S\_balanced = min(nFG / 7, nBG / 6)$$

where the numerators are unique supporting species. Requiring at least five FG and five BG species retained 2,984 events at 2,982 unique positions in 2,104 genes. Two parallel frozen branches were then distinguished. The sensitivity endpoint required FG dominant-amino-acid frequency ≥ 0.80 and retained 1,713 positions in 1,397 genes. The strict primary endpoint was restricted to primary events, required FG and BG support ≥ 5, complete FG concordance (dominant frequency = 1), and Patterns 1–2, and retained 1,068 positions in 934 genes: 600 Pattern-1 and 468 Pattern-2 events. An exploratory FG4/BG4 and FG-dominance ≥ 0.80 branch retained 18,959 unique positions in 7,264 genes. These branches were generated from the same hypothesis sample; multi-seed resampling and leave-one-species or leave-one-clade analyses were not performed and no robustness claim is based on them.

Unique-species support distributions and every frozen threshold endpoint are shown in supplementary figure S3; the strict and sensitivity event tables are supplementary data S3 and S4. A prespecified local-alignment audit would additionally require local gap fraction, FG/BG representation, low-complexity and call-clustering flags, coordinate and identifier agreement, paralogy or transcript warnings, and dated manual inspection of the aligned window. That audit has not been frozen. Representative alignments were therefore not selected, supplementary figure S4 was not produced, and the reported substitutions are candidates rather than functionally or alignment-validated residues.

#### Supplementary Method S4. External validation, functional enrichment, and sensitivity analyses

The common detectable universe for external validation and functional enrichment was the 16,133 gene identifiers represented by the screened alignments, including the 5,338 genes with header-only CAAStools output. Non-empty event files supplied their internal gene identifier; header-only files were parsed from their basenames. Identifiers were deduplicated before gene-level tests, and every query gene was required to occur in the background. The alignment identifiers were submitted as human gene symbols to g:Profiler; the service's returned mapping was retained rather than silently resolving failed or ambiguous names. This strategy submitted a biologically relevant detectable background instead of the whole human genome.

For external comparison, the unfiltered position table of Farré et al. (2021) was reprocessed using the published complete-discovery conditions: all six foreground and six background species present (`FFGN=6`, `FBGN=6`) and no foreground or background gap or missing species (`GFG=GBG=MFG=MBG=0`). These criteria retained 3,300 rows in 2,338 unique RefSeq accessions. Transcript version suffixes were removed before local mapping. UCSC hg38 and hg19 `refGene` tables dated 18 August 2020 were combined with hg38 `ncbiRefSeq` dated 13 August 2025 to reconcile publication-era and later symbols. All symbols were retained for accessions with multiple mappings, after which each tested set was deduplicated; a sensitivity analysis restricted the comparison to single-symbol accessions. In total, 2,334 of 2,338 accessions mapped to 2,365 symbols, of which 2,073 occurred in the 16,133-gene universe.

Agreement was tested separately for all Farré discovery scenarios, scenarios 1–2, and scenario 3. For each comparison, a one-sided hypergeometric over-representation test used N = 16,133 background genes, K detectable Farré genes, n = 934 strict CAAS genes, and x genes in their intersection. The exact probability was $P(X \geq x)$ under sampling without replacement. Observed overlap, expectation nK/N, observed/expected ratio, and exact nominal P value were archived. This analysis measures gene-level concordance only; it does not require recurrence of the same aligned position or amino-acid substitution. Complete mapping and overlap statistics are supplied in supplementary table S5.

Functional over-representation was evaluated with the official g:Profiler g:GOSt API (Reimand et al. 2007; Kolberg et al. 2023). The unordered query comprised 934 strict genes, and the custom background comprised all 16,133 screened genes (`organism=hsapiens`, `domain_scope=custom`). All annotation sources available to the service were included, as were electronically inferred Gene Ontology annotations; under-representation was not tested. Experiment-wide significance was defined as g:SCS-corrected $P \leq 0.05$. Two requests were retained: one returning all tested terms without evidence lookup and one returning significant terms with intersections and evidence. The API was queried on 26 August 2026 against database version `e114_eg62_p19_27110d83`. Of 934 submitted query genes, 917 mapped, three were flagged as ambiguous, and 14 failed; 28,930 terms were tested and 17 passed g:SCS correction. Request payloads, raw responses, mapping tables, failed identifiers, and complete term tables were archived in supplementary data S5, with significant terms summarized in supplementary table S6. Human Phenotype Ontology (HPO) memberships were treated as human annotation evidence, not as fertility measurements in the primates analysed.

The fertility evidence matrix was defined as the 31-gene union of the significant HPO terms Male infertility and Decreased fertility in males. For each gene, the matrix recorded membership in either HPO term, strict CAAS position and pattern, unique FG and BG support, and Farré scenario membership; sperm/flagellar classification was added for the sensitivity analysis. Alignment-quality status remained unassessed because the audit described in S3 was not frozen. Correlated HPO terms were not interpreted as independent replications. The complete matrix is provided in supplementary table S7.

Enrichment sensitivity was evaluated after two nested, prespecified exclusions. The strict set removed 12 cilia/flagella genes (`AKAP3`, `CFAP43`, `CFAP74`, `DNAH1`, `DNAH5`, `FSIP2`, `GAS2L2`, `LRRC56`, `NME8`, `RPGR`, `RSPH4A`, and `TTC12`), leaving a query of 922 genes. The extended set additionally removed `CYLC1`, `PMFBP1`, `SPATA16`, and `SUN5`, leaving 918 genes. The definitive analyses removed the same genes from query and background, producing backgrounds of 16,121 and 16,117 genes; 905 and 901 query genes mapped, respectively. Each reduced analysis repeated the original unordered g:Profiler request and g:SCS correction. As an independent verification, one-sided hypergeometric over-representation was recalculated after explicitly pruning each tested ontology term to the reduced universe; these manual probabilities were nominal and did not replace experiment-wide g:SCS inference. The exclusion design, mapping attrition, fertility-term intersections, and retention of significant terms are shown in supplementary figure S5 and reported completely in supplementary table S8. No Farré-overlap test was performed after these strict-set exclusions, so no claim is made that gene-level agreement with Farré et al. remains unchanged after sperm-structural gene removal.

#### Reproducibility and data files

Phenotype construction used Python 3.9.4 and pandas 2.2.3. Phylogenetic analyses used R 4.3.1, `ape` 5.8-1/5.8.1, `geiger` 2.0.11, and `phytools` 2.5.2. The molecular workflow used Python 3.11, OpenJDK 17, a frozen locally extended CAAStools 0.9 beta code bundle, and Nextflow DSL2; the production run identifier was `run-260819161121`. Fixed seeds were `20260817` for Blomberg's K permutations and `260811` for hypothesis selection. Source checksums are recorded in the stage-level audit files and each supplementary figure's `source.manifest.tsv`; plotting scripts read frozen source tables and validate their expected dimensions before rendering. Analysis coordinates are zero-based alignment columns, whereas species and gene identifiers preserve the frozen source conventions. A release URL/DOI and the exact Nextflow version were not present in the archived production record and must be added to the final data-availability package before submission.

Archived inputs comprise the phenotype source tables, the Kuderna tree, the protein-alignment collection, and the Farré comparison table. Analysis outputs comprise the audited phenotype table, pairwise PSS table, run-level hypothesis manifest, pooled event tables, enrichment responses, and overlap summaries. Each supplementary-figure directory separately contains plot-specific data, a source manifest with checksums, the rendering script, session information, and the PDF, SVG, PNG, and TIFF publication artifacts. These layers were kept distinct so that figure builders could validate and display frozen results without silently recalculating or filtering the primary analyses.

### Supplementary Figure Legends

#### Supplementary Figure S1. Phenotype provenance and diagnostics for the primate longevity quotient

**A,** Inclusion and reconciliation flow derived from the frozen phenotype audit. Of 526 source records, 364 lacked a finite positive maximum lifespan, body mass, or both. The remaining 162 records had normalized species names and valid traits; 36 were absent from the Kuderna S4 phylogeny under exact-name reconciliation, leaving 126 species in the analysis. **B,** Observed maximum lifespan against the fixed mammal-wide allometric expectation, calculated as 6.47 × body mass in grams$^{0.189}$. The dashed diagonal denotes equality between observed and expected lifespan. Foreground (FG) species are shown as filled vermillion circles, background (BG) species as outlined blue triangles, and other species as grey circles. **C,** Ordered distribution of the untransformed longevity quotient (LQ = observed/expected maximum lifespan) for all 126 analysed species. The central dashed line marks LQ = 1; blue and vermillion dashed lines mark the operational BG and FG thresholds of 0.8 and 1.3. The seven FG and six BG species are labelled and connected to their positions. Thresholds are classification rules rather than confidence limits. **D,** Family-level counts of analysed records, trait-valid records not matched to the phylogeny, and records excluded for missing or non-positive traits. Counts diagnose taxonomic coverage and are not tests of lineage differences. Family names are shown as supplied by the frozen audit.

#### Supplementary Figure S2. Complete phenotype-shift distribution and construction of the FG and BG species sets

**A,** All 7,875 unique unordered species pairs ranked by phenotype-shift score (PSS). The deterministic upper and lower 1% tails each contain 79 pairs; the high-PSS tail is shown in vermillion and the low-PSS tail in blue. Tail membership is descriptive and is not a significance test. **B,** Number of upper- and lower-tail pairs containing each species represented in either tail. Symbol size distinguishes species appearing in both tails from species appearing in only one tail. The 47 species present in both tails are grey unless they meet the final longevity-quotient threshold; the seven FG species are filled vermillion circles and the six BG species are outlined blue triangles. Pair counts do not constitute independent biological replicates. **C,** Ordered LQ values for the 47 shared-tail species. Fixed dashed lines mark LQ = 0.8 and LQ = 1.3, and the central dashed line marks LQ = 1. Applying strict inequalities within this candidate set yields seven FG species with LQ > 1.3, six BG species with LQ < 0.8, and 34 intermediate species. **D,** Complete 126-tip Kuderna S4 tree pruned to the phenotype analysis set. Outer tracks, from inner to outer, encode LQ magnitude, upper-tail membership, lower-tail membership, and final FG/BG status. Branches show phylogenetic placement only; no ancestral states or longevity shifts are inferred.

#### Supplementary Figure S3. Species-level support and filtering sensitivity of the pooled CAAS discovery

**A,** Exact distribution of balanced unique-species support, defined as min(FG support/7, BG support/6), across all 129,622 nominally significant pooled events. The vertical dashed line marks the balance implied by a minimum of five FG and five BG species. Counts are drawn from the frozen exact-frequency table and shown on a logarithmic axis. **B,** Joint counts of unique FG and BG support in the same broad event universe. Outlined cells satisfy FG ≥ 5 and BG ≥ 5 and contain 2,984 events in 2,104 genes. Repeated hypotheses do not contribute additional support. **C,** Auditable grid of all available frozen endpoints. The nominal universe contains 126,598 unique gene-position combinations in 10,795 genes. The exploratory FG4/BG4, FG-dominance ≥ 0.80 endpoint contains 18,959 positions in 7,264 genes. The FG5/BG5 support filter contains 2,982 positions in 2,104 genes. Requiring FG-dominance ≥ 0.80 retains 1,713 positions in 1,397 genes. The primary strict endpoint requires FG5/BG5 support, complete FG amino-acid concordance, and Patterns 1-2, retaining 1,068 positions in 934 genes. The FG4/BG4 row is exploratory; the 1,713-position row is a parallel sensitivity analysis. **D,** Joint FG/BG support within the strict set, separated by pattern. Pattern 1 contributes 600 positions and Pattern 2 contributes 468. Each position contributes once to its unique-species support combination. CAAS, convergent amino-acid substitution; FG, foreground; BG, background.

#### Supplementary Figure S5. Dependence of functional enrichment on sperm cilia and flagella structural genes

**A,** Prespecified query-and-background sensitivity design. The original analysis submitted 934 strict CAAS genes against a 16,133-gene background; 917 genes mapped and 17 terms were significant. Removing 12 strict cilia/flagella genes from both query and background produced 922 submitted genes, 905 mapped genes, a 16,121-gene background, and 15 significant terms. Extending the exclusion to 16 sperm-structural genes produced 918 submitted genes, 901 mapped genes, a 16,117-gene background, and 15 significant terms. **B,** g:SCS-corrected significance for the two male-fertility HPO terms, Adult onset, and representative retained non-fertility terms. Points at zero correspond to corrected $P = 1$; the vertical dashed line marks corrected $P = 0.05$. Male infertility changed from corrected $P = 3.43 \times 10^{-4}$ to 1 after both exclusions, and decreased fertility in males changed from $7.94 \times 10^{-4}$ to 1. Adult onset also lost significance, whereas the displayed extracellular-matrix, actin-projection, microvillus, and brush-border terms remained significant. **C,** Fertility-term intersection sizes. Male infertility declined from 29 genes to 17 after strict exclusion and 13 after extended exclusion; decreased fertility in males declined from 31 to 19 and 15 genes, respectively. Loss of experiment-wide significance is distinct from biological disappearance of the remaining annotated genes. **D,** Significance matrix for the 18-term union and membership matrix for the excluded genes. Fourteen original terms remain significant in all analyses; the two fertility terms and Adult onset lose significance, whereas one kidney proximal-tubule microvillus term becomes significant after both exclusions. Thus, 15 terms are significant under both exclusions. The strict 12-gene set is nested within the extended 16-gene set. HPO annotations are functional annotations rather than measured fertility phenotypes in the analysed primates. CAAS, convergent amino-acid substitution; HPO, Human Phenotype Ontology.

### Supplementary Figure S1

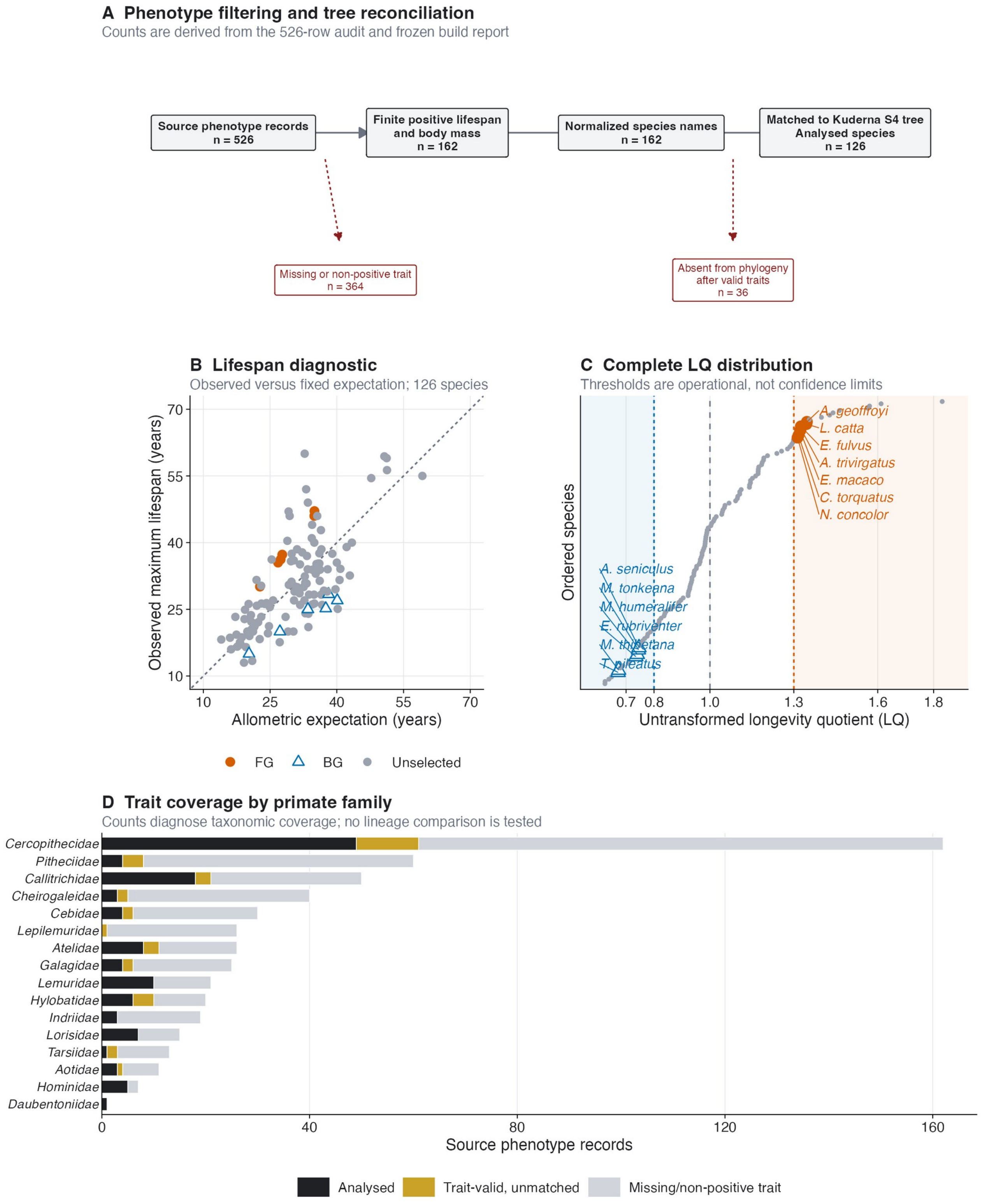

#### Supplementary Figure S2

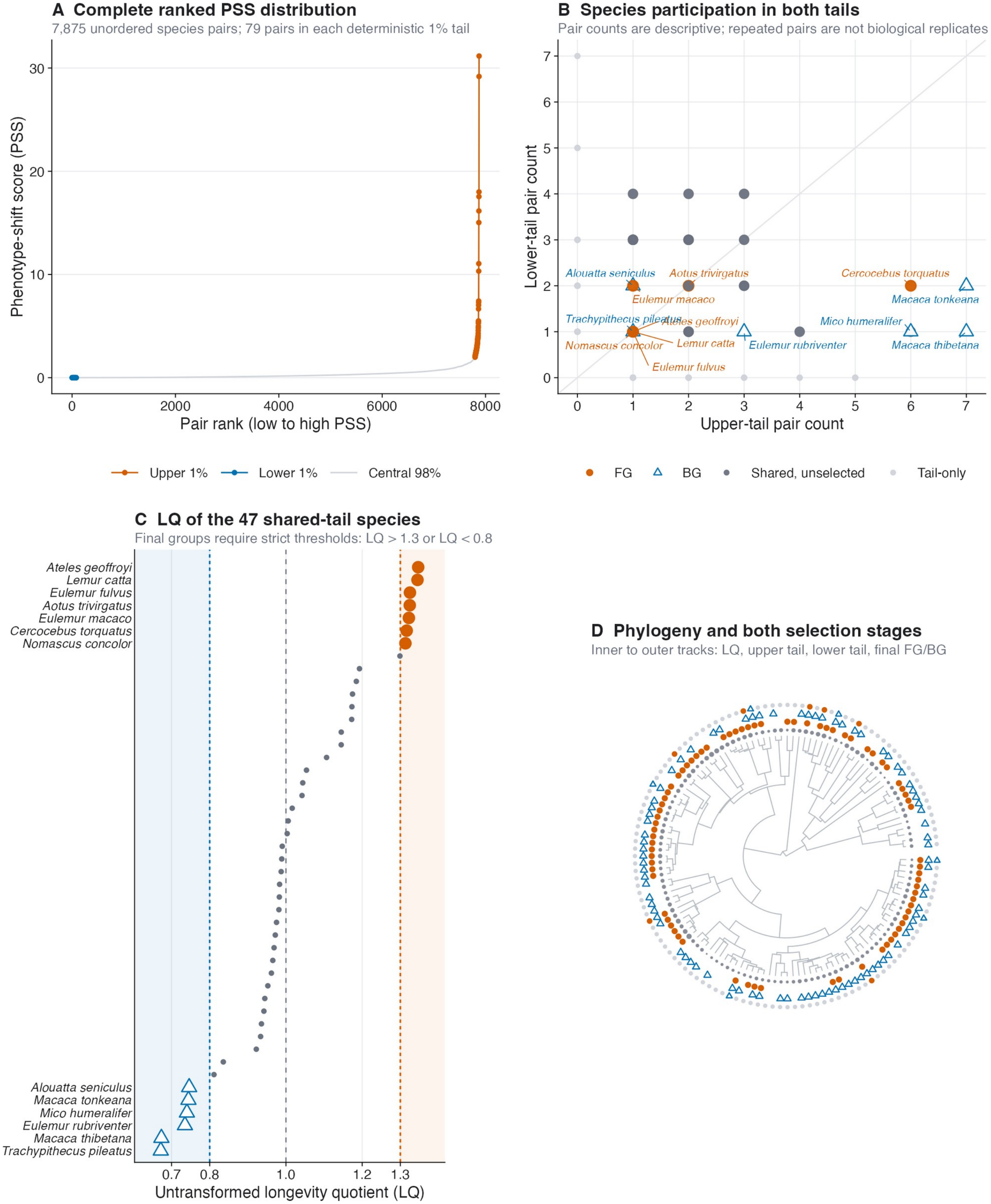

### Supplementary Figure S3

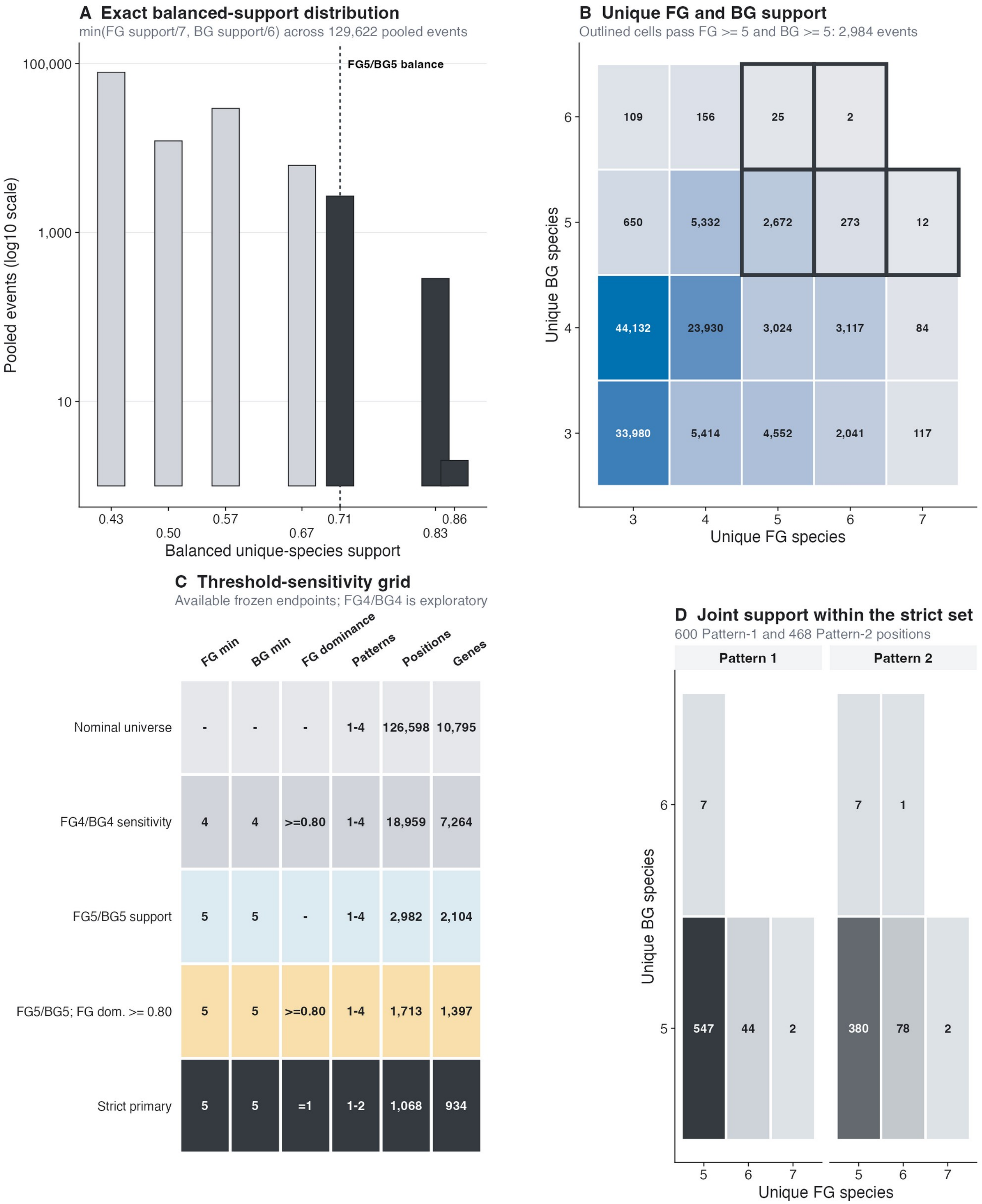

### Supplementary Figure S5

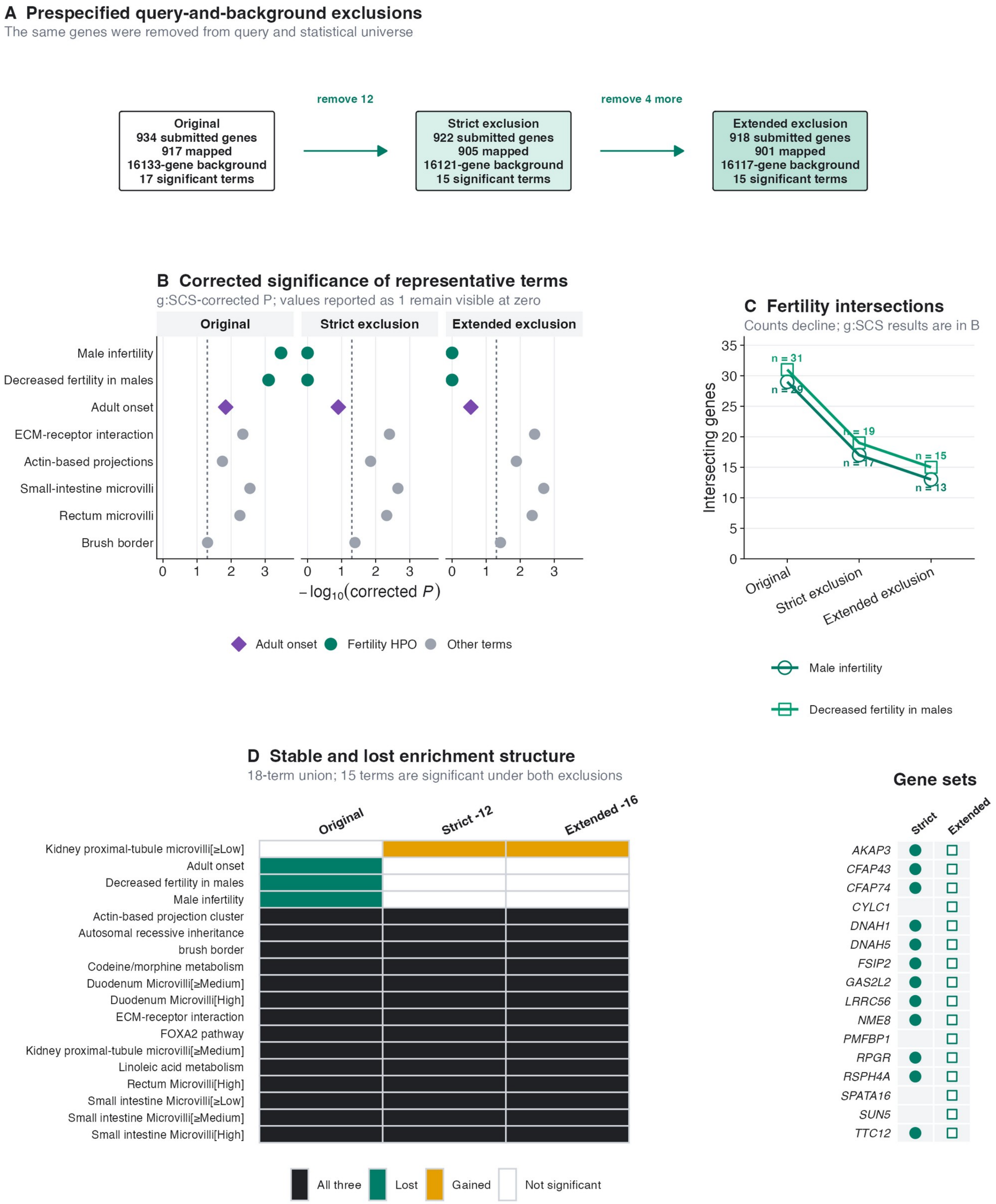
